# Unsupervised Clusterless Decoding using a Switching Poisson Hidden Markov Model

**DOI:** 10.1101/760470

**Authors:** Etienne Ackermann, Caleb T. Kemere, John P. Cunningham

**Affiliations:** Department of Electrical and Computer Engineering, Rice University, Houston, TX 77005-1892, USA; Department of Statistics, Columbia University, New York, NY 10027, USA

**Keywords:** Hidden Markov Models, Clusterless Decoding

## Abstract

Spike sorting is a standard preprocessing step to obtain ensembles of single unit data from multiunit, multichannel recordings in neuroscience. However, more recently, some researchers have started doing analyses directly on the unsorted data. Here we present a new computational model that is an extension of the standard (unsupervised) switching Poisson hidden Markov model (where observations are time-binned spike counts from each of *N* neurons), to a clusterless approximation in which we observe only a *d*-dimensional mark for each spike. Such an unsupervised yet clusterless approach has the potential to incorporate more information than is typically available from spike-sorted approaches, and to uncover temporal structure in neural data without access to behavioral correlates. We show that our approach can recover model parameters from simulated data, and that it can uncover task-relevant structure from real neural data.

## 1. Introduction

Decoding neural activity is fundamental to much of neural data analysis. Most of the approaches to decoding are *supervised*, meaning that we are given labeled data (e.g., a set of an animal’s positions along a track), as well as the corresponding neural data, and our task becomes to learn a mapping from the neural activity to position, or whichever other extrinsic correlate we are interested in. In this context, neural activity typically refers to spike events from an ensemble (tens, hundreds or maybe even thousands) of neurons. However, neural data are most commonly recorded as continuous voltage traces on multiple electrode channels, so that prior to decoding, spikes first have to be detected from these voltage traces, and each of those spikes has to be assigned to one of *N* neurons in a process called *spike sorting* (Gerstein and Clark, 1964; Lewicki, 1998). Figure 1.A shows an example of the information that may be available for supervised decoding with spike sorted data, including spike timing information, the neuron identity for each spike, spike waveform feature information, and the extrinsic correlates (position in this figure).

**Figure 1:**
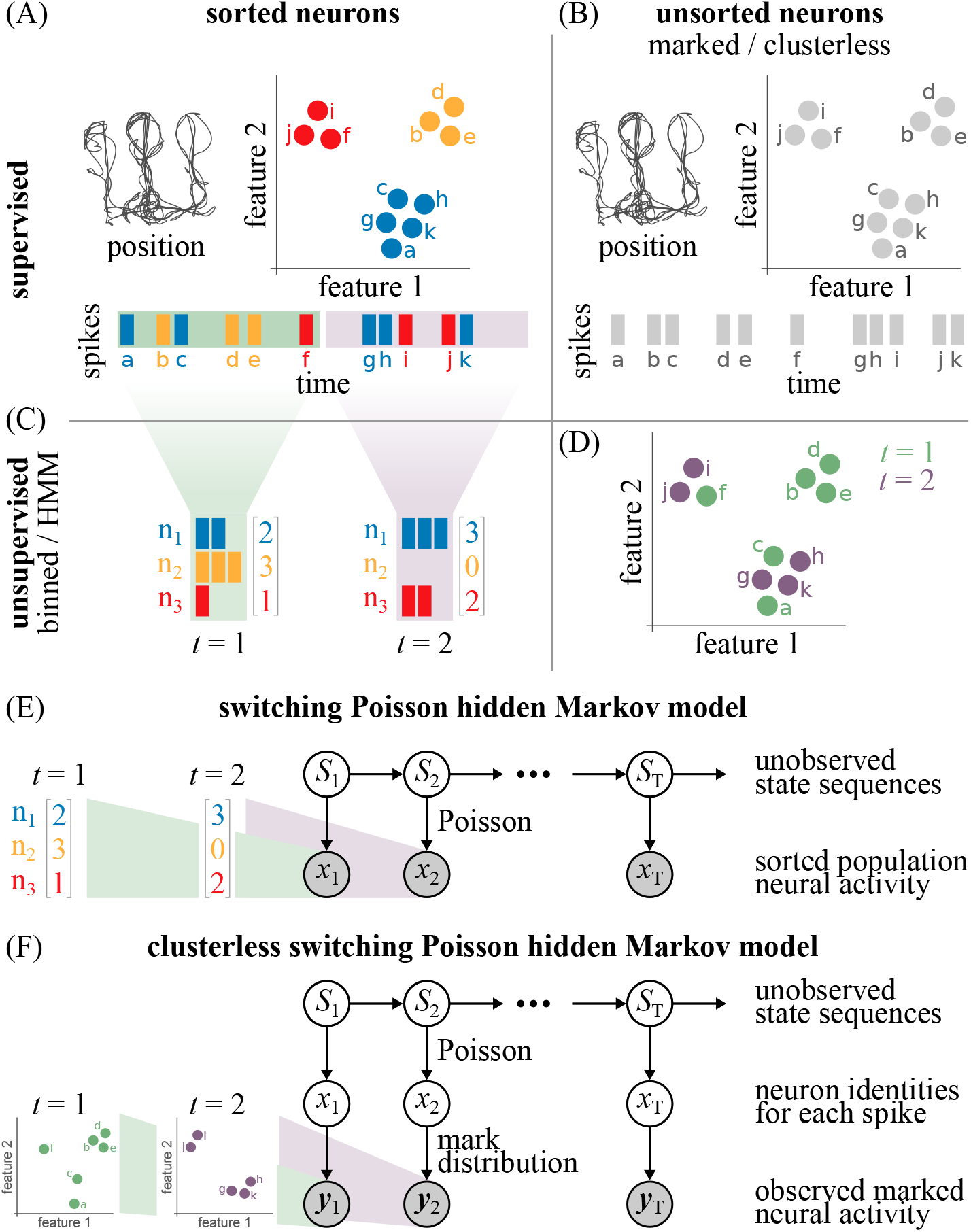
Switching Poisson HMM versus clusterless HMM. (A)-(D) Illustration of information available within different decoding paradigms. (A) Supervised + sorted decoding. (B) Supervised + clusterless decoding. (C) Unsupervised + sorted decoding (e.g., switching Poisson HMM). (D) Unsupervised + clusterless decoding (this paper). (E) Graphical model illustrating the switching Poisson HMM, where the sequence of states are unobserved, and each state generates an observable collection of spike counts within the associated time window. Examples of observations during two time windows are shown on the left, namely spike counts with neuron identities. (F) Graphical model for our clusterless HMM, where the neuron identities generating the spikes are unobservable. The unobservable neuron identities become latent variables in our model, and these generate the observable marks. Examples of observations are shown on the left, namely a collection of unsorted spike waveform features during each time window.

Spike sorting workflows are often characterized by a high degree of subjectivity and variability (Febinger et al., 2018), and even with fully automated spike sorting approaches like that developed by Chung et al. (2017), results may still be sensitive to a host of parameter choices. Spike sorting also throws away potentially valuable information: most spikes are discarded in the spike sorting process for being too small or too “difficult” to sort, and so they get lumped into background multiunit activity or “hash”. For those spikes that are sorted, we typically throw away the uncertainty information about each spike assignment. With the rapid rise in popularity of data mining and machine learning, it is not surprising that researchers have tried to find ways to interpret neural data directly, without having to perform spike sorting first. Indeed, the desire to circumvent the subjectivity and tedium of the spike sorting process, along with the observation that spike sorting throws away valuable information, has led to the recent development of many so-called “clusterless” approaches (Ventura, 2008; Kloosterman et al., 2013; Deng et al., 2015). Clusterless approaches operate directly on spike waveform features (such as the peak amplitudes on each of the four channels of a tetrode), without requiring or assuming knowledge of the underlying neuron identity that generated each individual spike. Figure 1.B illustrates typical information that may be available during supervised clusterless decoding, namely spike timing information, spike waveform features, and the behavioral correlates.

A complimentary paradigm to the supervised decoding approaches mentioned above, is that of *unsupervised* approaches to understand and interpret neural data (see e.g., Low et al., 2018; Williams et al., 2018; Maboudi et al., 2018; Williams et al., 2019; Mackevicius et al., 2019). These unsupervised approaches provide powerful ways of understanding the latent dynamics of neural activity, but to date they all require sorted spikes from ensembles of neurons. Figure 1.C shows an example of the information available during unsupervised decoding of sorted data, namely the spike timing information and the neuron identities responsible for generating those spikes. Note that there are no (observable) extrinsic correlates, and that fine spike timing information is lost during the binning process.

In this paper, we extend one particular unsupervised approach, namely the (switching Poisson) hidden Markov model (HMM; see e.g., Linderman et al., 2016; Maboudi et al., 2018, as well as Figure 1.E) to the clusterless setting, in a new model that we call the *clusterless* hidden Markov model. This clusterless HMM builds on the switching Poisson framework used by Gat and Tishby (1993); Kemere et al. (2008), and others, by treating the neuron identities responsible for the spikes as latent variables in the model (see Figure 2.F), and by incorporating our uncertainty about those identities in the form of parametric distributions of the waveform features associated with each hidden neuron, the *mark distributions*. Such a clusterless approach allows us (i) to incorporate more information than typically available with a standard spike sorting approach, and (ii) to eliminate the subjectivity and variability arising from the spike sorting process. The incorporation of additional information is particularly appealing for short time-scale event analysis (e.g., replay or theta sequences), where there are often not sufficiently many spikes remaining after spike sorting to decode with much accuracy.

**Figure 2:**
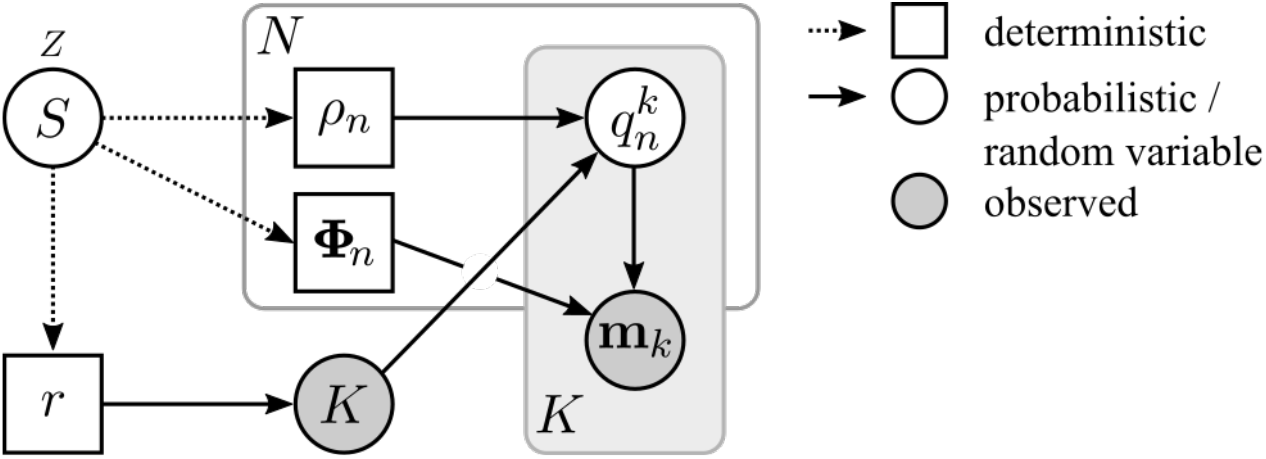
Probabilistic graphical model during single time window for the clusterless HMM. *S*: discrete state, |*S*| = *Z*; *r*: aggregate firing rate, *ρ_n_*: relative firing rate of neuron *n*; *K* ~ Pois(*r*): number of marks observed; *q_k_* ~ Multinom(*K*, ***ρ***): neuron identity indicator; **Φ**_*n*_: mark distribution parameters; **m**_*k*_: *d*-dimensional mark.

We present the clusterless HMM and one possible way of doing inference on the model in Section 2, and we show that we can accurately recover the underlying model parameters in a simulation study in Section 3. We also show that the model can uncover temporal structure in real neural data from the rat hippocampus in Section 4, and finally we provide some directions for future improvement and investigation in Section 5.

## 2. Methods

We present a novel clusterless HMM for analyzing and decoding multiunit sequential neural activity in an unsupervised manner. The Baum–Welch algorithm is an expectation maximization (EM)-based algorithm for estimating the maximum-likelihood parameters of an HMM (Bilmes et al., 1998), and we will use it in this paper to do inference on our model. The Baum–Welch algorithm critically relies on our ability to evaluate *P*(**y**_*t*_ | *S_t_*), from which we can efficiently compute several other quantities of interest to enable inference with our models (see e.g., Rabiner (1989) and Appendix B). That is, we need to be able to evaluate the probability of observing a particular outcome **y**_*t*_ for each (hidden) state *S_t_*. In the following sections, we will present our full clusterless HMM, and develop a sampling-based approach to approximate *P*(**y**_*t*_ | *S_t_*).

The idea here is very simple. In particular, *if we did know the neuron identities for each spike*, then we would not have to use a “clusterless” approach, and we could instead fit a standard switching Poisson HMM. So in our clusterless model, we can similarly compute probabilities *if we assume that we know the neuron identities*, and we simply treat these identities as hidden or latent variables in our model. Moreover, we do not need to know exactly how many neurons there are, since a specification of the number of neurons simply determines the partitioning of our waveform feature space, so that an over-specification (or perhaps even a small under-specification) of the number of neurons should not have a significant effect on our model’s behavior. These ideas will be made precise below.

### 2.1 Notation and preliminaries

Suppose that we record from a population of *N* neurons, with each neuron identified by *n* ∈ {1, 2, … *N*}. Further, suppose that we observe a *d*-dimensional mark 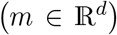 for each spike from a neuron. The marks could be, for example, the peak amplitudes from the four channels of a tetrode, or principal components of the observed waveform on a collection of electrodes, and so on.

If we assume that the neurons have *state*-dependent firing rates 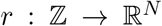 with *S_t_* ↦ *r*(*S_t_*) ≡ **r**_*t*_, then we consider each neuron as generating a train of events from an inhomogeneous (state-dependent) Poisson process, with an associated (state-dependent) rate, 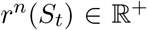. To be more precise, we assume that the neurons are *independent*, and that they have *N* state-dependent rates, **r**_*t*_ = (*r*^1^*S_t_*), *r*^2^(*S_t_*), …, *r^N^*(*S_t_*)) at time 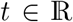, and for some state *S_t_* ∈ {1, 2, …, *Z*}. We may then collect all of these rates as the matrix 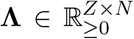, and we will estimate these rates as part of the HMM parameter estimation process.

For simplicity, we will assume that we have only one probe (or tetrode). In general, we assume that the tetrodes are independent, so that tetrodes record from disjoint subsets of neurons, and therefore generalization to the multiprobe case is straightforward.

Now, consider a single observation window (*t* – 1,*t*] simply identified with *t*, during which we observe *K*(*t*) = *K* spike events (equivalently, we observe *K* marks: **y**_*t*_ = {*m*_1_, …, *m_K_*}, 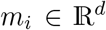, *i* = 1, …, *K*), and for which we assume that each neuron has a constant firing rate for the duration of the time window. Since observation windows are conditionally independent (given the underlying states), we only need to concern ourselves with evaluating *P*(**y**_*t*_ | *S_t_*) for a single observation window (the process remains unchanged for each observation window).

Finally, let us assume that the marks (the spike waveform features) can be modeled as coming from unit-specific distributions parameterized by **Φ**. The choice of this distribution will depend on which features are ultimately chosen to represent each mark. In this paper, we simply use the peak amplitudes on each of the tetrode channels, which can be adequately modeled as coming from unit-specific multivariate normal distributions in practice.

### 2.2 Model Specification

As suggested before, let **y**_*t*_ denote the *observation* at time *t*, where 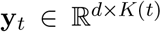, and where *K*(*t*) is the number of marks observed during time window *t*. We assume that the observations are sampled at discrete, equally-spaced time intervals, so that *t* can be an integer-valued index.

The hidden state space is assumed to be *discrete*, with *S_t_* ∈ {1, …, *Z*}. To define a probability distribution over sequences of observations, we need to specify a probability distribution over the initial state, **π** ≡ *P*(*S*_1_), with **π_*i*_** = *P*(*S*_1_ = 1), the *Z* × *Z* state transition probability matrix, **A**, with *a_ij_* = *P*(*S_t_* = *j* | *S*_*t*–1_ = *i*), and the emission model defining *P*(**y**_*t*_ | *S_t_*).

In the usual multiple-spike-train switching Poisson model (see e.g., Jones et al., 2007; Kemere et al., 2008), we simultaneously observe *N* spike trains (corresponding to *N* neurons). Each spike train is modeled as an independent Poisson process with rate λ_*n*_(*t*), and the vector of firing rates, **λ**(*t*), switches randomly between one of *Z* states. We will use the same idea here, but since the number of neurons and more importantly the identities of those neurons are *hidden*, we will have to be slightly more careful in our treatment. Nevertheless, assuming that our data were generated by *N* hidden neurons, we associate the N-dimensional rate vector **r**_*t*_ with the expected number of marks from each of the neurons during time window *t*. There are *Z* possible rate vectors (one for each state *S_t_*), so that we can collect all the possible rates captured by the model in the rate (or observation) matrix 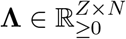.

We further assume that our model is time-invariant in that the state transition probability matrix and the emission model do not change over time, and that the states form a first-order Markov chain, namely that *P*(*S_t_* | *S*_*t*–1_, …, *S*_1_) = *P*(*S_t_* | *S*_*t*–1_). The HMM is then fully specified by the set of parameters **Θ** = **{π, A, Λ}**. There are therefore a total of *Z*(1 + *Z* + *N*) parameters to estimate for the model^1^.

### 2.3 Sampling approach to evaluate *P*(y_*t*_ | *S_t_*)

In order to use the Baum-Welch algorithm to estimate the set of parameters **Θ** = **{π, A, Λ}**, we need to be able to evaluate

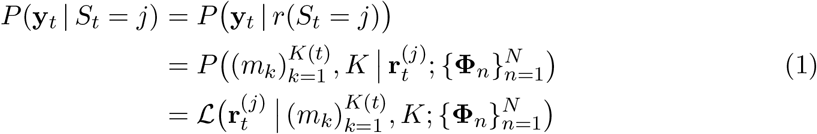

where the dependence of the sequence of observed marks 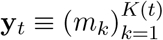 on the hidden state *S_t_* is realized through the state-dependent firing rates 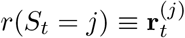, as well as the unit-specific mark distribution parameters 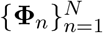. That is, conditioning on *S_t_* is equivalent to conditioning on 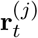. Note also the 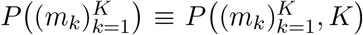, and we will simply write 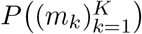 for convenience.

A graphical representation during one time window of our clusterless HMM is shown in Figure 2. The collection of *K* marks depend on the unobserved neuron identities 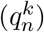, as well as on the possibly state dependent mark distribution parameters, **Φ**. The neuron identities depend on the firing rates, *ρ* and *r*, and on how many marks were observed (*K*). The number of marks, *K*, is assumed to follow a Poisson distribution with mean rate *r*, the total number of expected spikes from all the neurons in state *S*.

Dropping the dependence of *K* on *t* (purely for notational simplicity), and dropping the explicit parameterization of *P*(·) by **Φ**, we note that we desire to evaluate

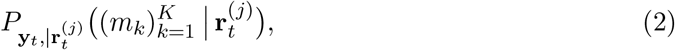

which is hard to evaluate explicitly for two key reasons, namely (i) we do not know which neurons / units gave rise to each of the marks, and (ii) we do not know *a priori* how many marks we will observe in a particular observation window, so that we cannot easily specify a simple (possibly multivariate) probability distribution over the sequence of observed marks. Indeed, the dimensionality of this distribution will have to depend on how many marks we observe in each window.

To address challenge (i) above, we introduce the auxiliary hidden random variable 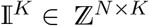, that encodes which of the latent units gave rise to each of the observed marks. Each column of 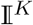 is a standard unit vector **e**_*n*_ whose elements are all zeros, except for the *n*th element, which is equal to one. That is, the *k*th column encodes the neuron identity *n* that generated the *k*th mark. We denote this as 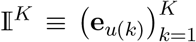, such that 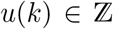 is the neuron identity of the *k*th mark.

In particular, we note that

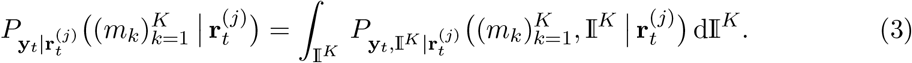

Making the dependence on time *t* and state *j* implicit (i.e., letting **r** ≡ *r*(*S_t_* = *j*) as before), we note that

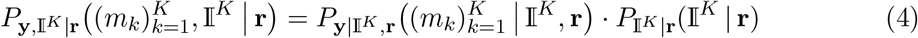

so that

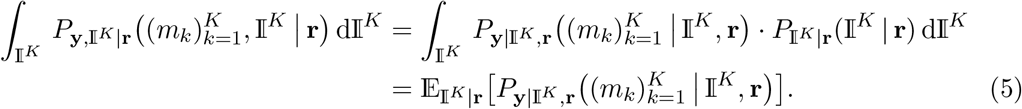

Note that it is generally difficult to compute the integral directly, whereas it is somewhat simpler to estimate the expected value, assuming of course, that we can sample from 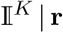 according to it’s distribution. Indeed, if we are able to sample 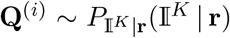, where 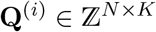 for *i* = 1, …, *M*, then

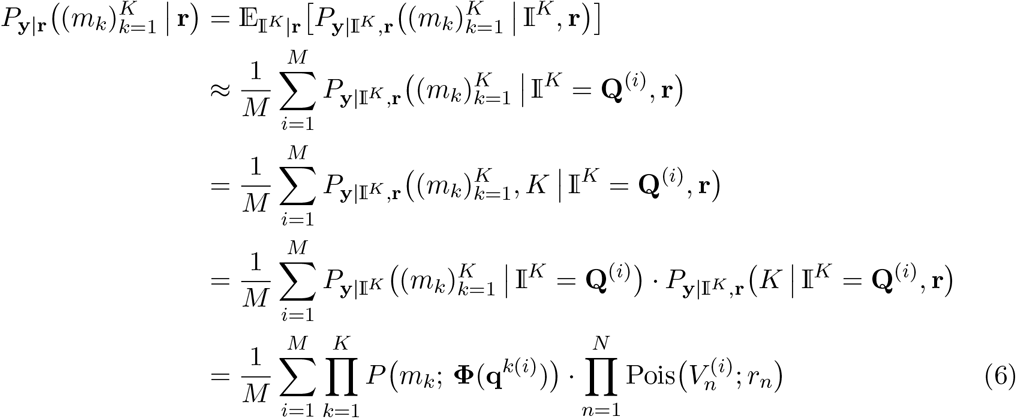

where we have explicitly brought *K* back into 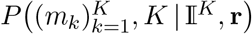, and where we have used the fact that the marks are assumed independent to express 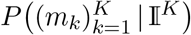 as a product. We have also made use of the conditional independence (see Figure 2) of the marks on the neuron identities (**Q**) and the number of observed marks (*K*) to factorize 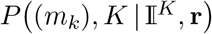. Then we can finally approximate the data log likelihood simply as:

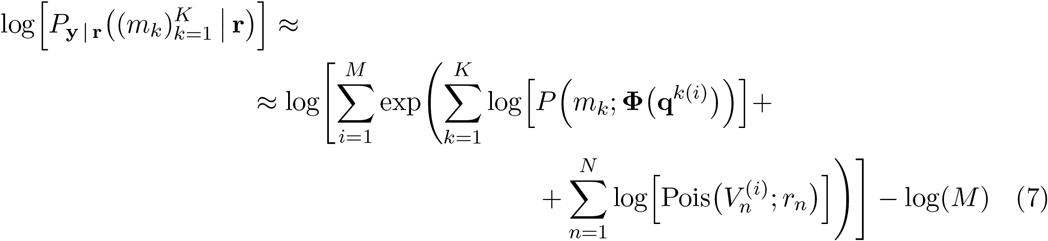

where, for each mark, **Φ** depends on the neuron identity encoded by **q**^*k*(*i*)^, the *k*th column of the *i*th sample **Q**^(*i*)^, and where 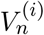 is the total number of spikes from neuron *n*, for the *i*th sample. That is,

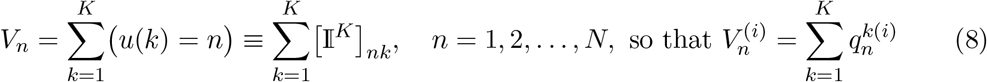

and where 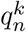 is the *n*th element of the *k*th column of **Q**, which equals one if we assume that the *k*th mark was generated by the *n*th neuron, and zero otherwise.

In order to sample 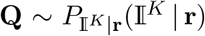, it is convenient to factorize the rate (**r**) into the aggregate rate 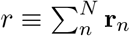, and the relative rates 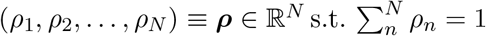 (see Figure 2). In this way, the (true) rate associated with neuron *n* is simply **r**_n_ = *rρ_n_*, and we can simply sample from the multinomial distribution of **V**, one trial at a time, for each mark identity in **Q**^(*i*)^. That is, for *k* = 1, …, *K*, sample 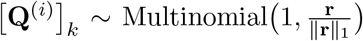, where [·]_*k*_ is used to denote the *k*th column:

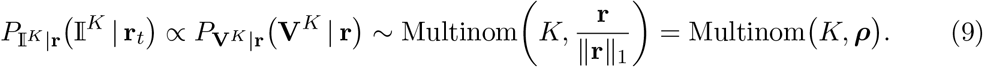

Note that this sampling strategy yields the correct number of neuron identities (proportional to their rates), but it does not affect the order of the columns of **Q**.

### 2.4 Updating the state-dependent firing rates, r^(*j*)^

Ordinarily, we may consider updating the rates according to

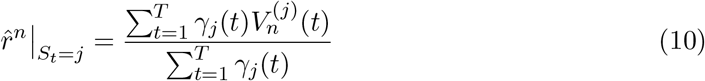

where 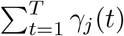 is the expected number of times that we are in state *j* (or equivalently, the expected number of transitions away from state *j*; even more explicitly, *γ_j_* (*t*) = *P*(*S_t_* = *j* | **Y**)), and 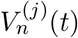 is the number of marks in time window *t* that were generated / emitted by neuron *n* (and given that the system is currently in state *j*). However, we do not know which marks were generated by which neurons, so we have to consider

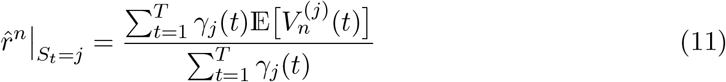

instead.

It turns out that we can efficiently compute this expectation, for time window (*t* – 1, *t*], as follows:

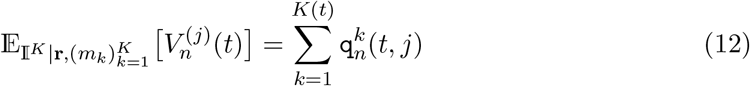

where

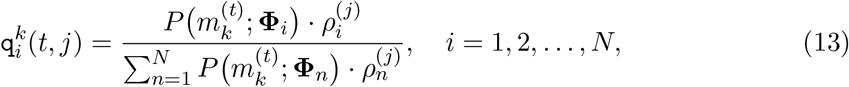

and where the dependence on *t* should be clear, namely that the sequence of marks (*m_k_*) are those observed during time window *t*. Note that (13) is computed *independent* of any sampling.

The prior distribution over states, **π_i_** ≡ *P*(*S*_1_ = *i*), and the transition probability matrix, **A**, can be estimated in the standard way, namely

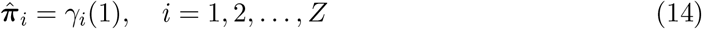

and

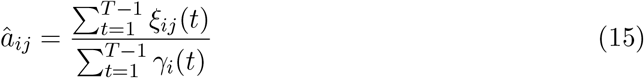

where *ξ_ij_*(*t*) = *p*(*S_t_* = *i*, *S*_*t*+1_ = *j* | **Y**), so that 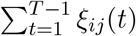 is the expected number of transitions from state *i* to state *j*.

The calculations of *P*(**y**_*t*_ | *S_t_*) and 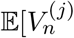 as well as the parameter estimation process are summarized in Algorithms 3–1 in Appendix B.

## 3. Simulation Case Study

We tested our clusterless HMM on simulated data so that we could determine exactly how well model parameters were recovered. In particular, we were interested in recovering the state transition probability matrix 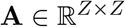, the observation—or rate—matrix 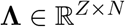, and a sequence of latent states (*S*_1_, *S*_2_, … *S_T_*).

For the remainder of this paper, we will assume that the mark distributions are independent of the hidden states, and that the marks can be modeled as multivariate normal distributions with parameters 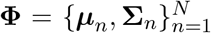 (see Figure 3). In this case, we can estimate the mark distributions before doing parameter estimation for our clusterless HMM, and we chode to use a simple Gaussian mixture model (GMM) to estimate the mark distribution parameters. Moreover, substituting the multivariate normal mark distributions into (6), (7), and (13) lead to

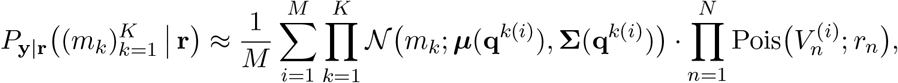

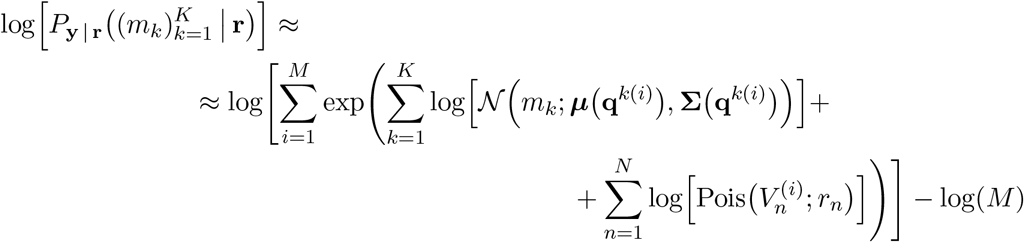

and

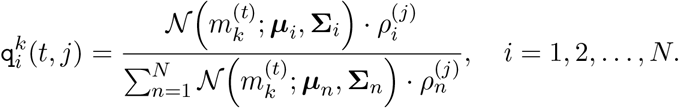

**Figure 3:**
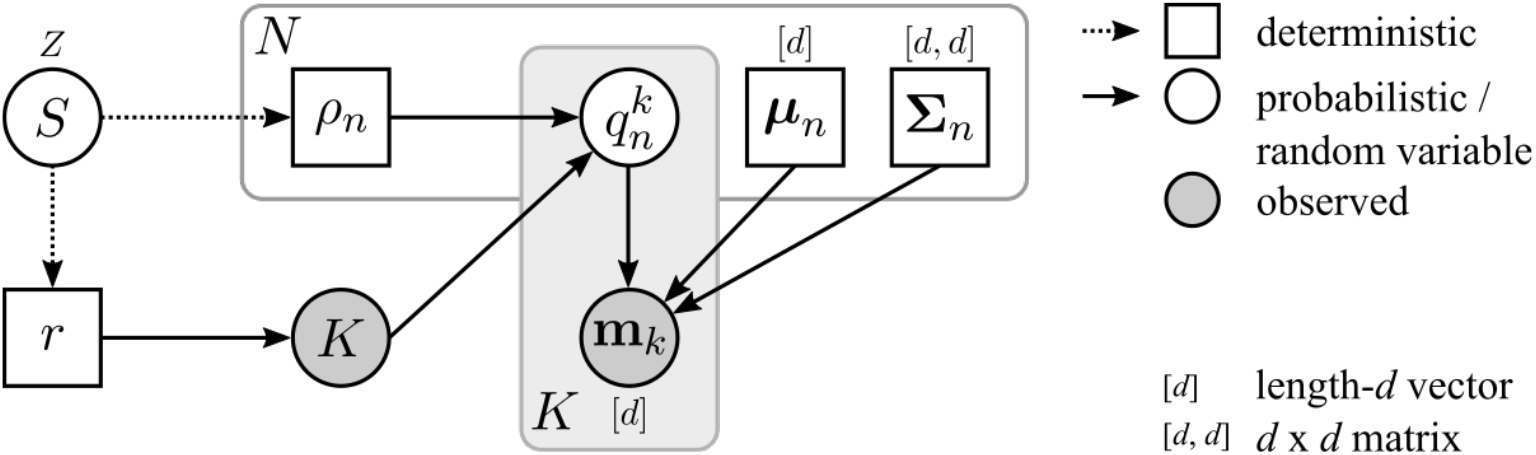
Probabilistic graphical model for the clusterless HMM with state-independent multivariate normal mark distributions. *S*: discrete state, |*S*| = *Z*; *r*: aggregate firing rate, *ρ_n_*: relative firing rate of neuron *n*; *K* ~ Pois(*r*): number of marks observed; *q_k_* ~ Multinom(*K*, ***ρ***): neuron identity indicator; ***μ***_*n*_: neuron centroid; **Σ**_*n*_: neuron covariance; 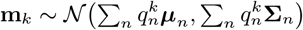.

### 3.1 Simple two-state, three-neuron example

As a simple yet representative example, we present parameter estimation results from a system with state transition probability matrix 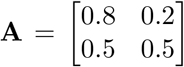 and a rate matrix 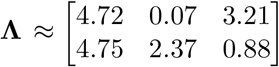.

More specifically, we randomly sampled the centroids and covariances of our *N* = 3 latent units, which we modeled as (*d* = 2 dimensional) multivariate normal distributions. This choice of mark distribution works reasonably well in practice when the marks are waveform peak amplitudes on channels of a tetrode, and we can use a Gaussian mixture model as a preprocessing step to estimate the parameters **Φ**_*n*_ = (***μ**_n_*, **Σ**_*n*_) for each neuron *n* = 1, …, *N*. In the rest of this paper we will adopt this strategy. Consequently, we now have that 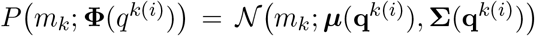 in (6) and (7), and that 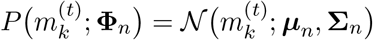 in (13).

We then sampled a sequence of states (*S*_1_, *S*_2_, …, *S_T_*) from the state transition probability matrix, **A**, and for each of those states we generated state-dependent spike events by sampling from a multivariate Poisson distribution with rates given by the corresponding row of **Λ**. For each of those spike events, we then generated a *d* = 2 dimensional mark by sampling from the unit-specific multivariate normal distributions, leading to a sequence of observations (**y**_1_, **y**_2_, …, **y**_*T*_).

Using the sequence of observations, 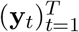, we then fit the clusterless HMM by evaluating *P*(**y**_*t*_ | *S_t_*) using (6) for each possible state *S_t_* and time window *t*, and by using (15) to update the transition probability matrix, **A**, and (11) to update the observation / rate matrix, **Λ**, during each iteration of the Baum-Welch algorithm. We used *M* = 5000 samples to approximate (6).

We sampled a length *T* = 200 state sequence, and used the first 100 samples to fit the model, and the next 100 samples to evaluate the Viterbi (state-) decoding accuracy of the model. The results from the parameter estimation process, as well as from the state decoding process, are shown in Figure 4.

**Figure 4:**
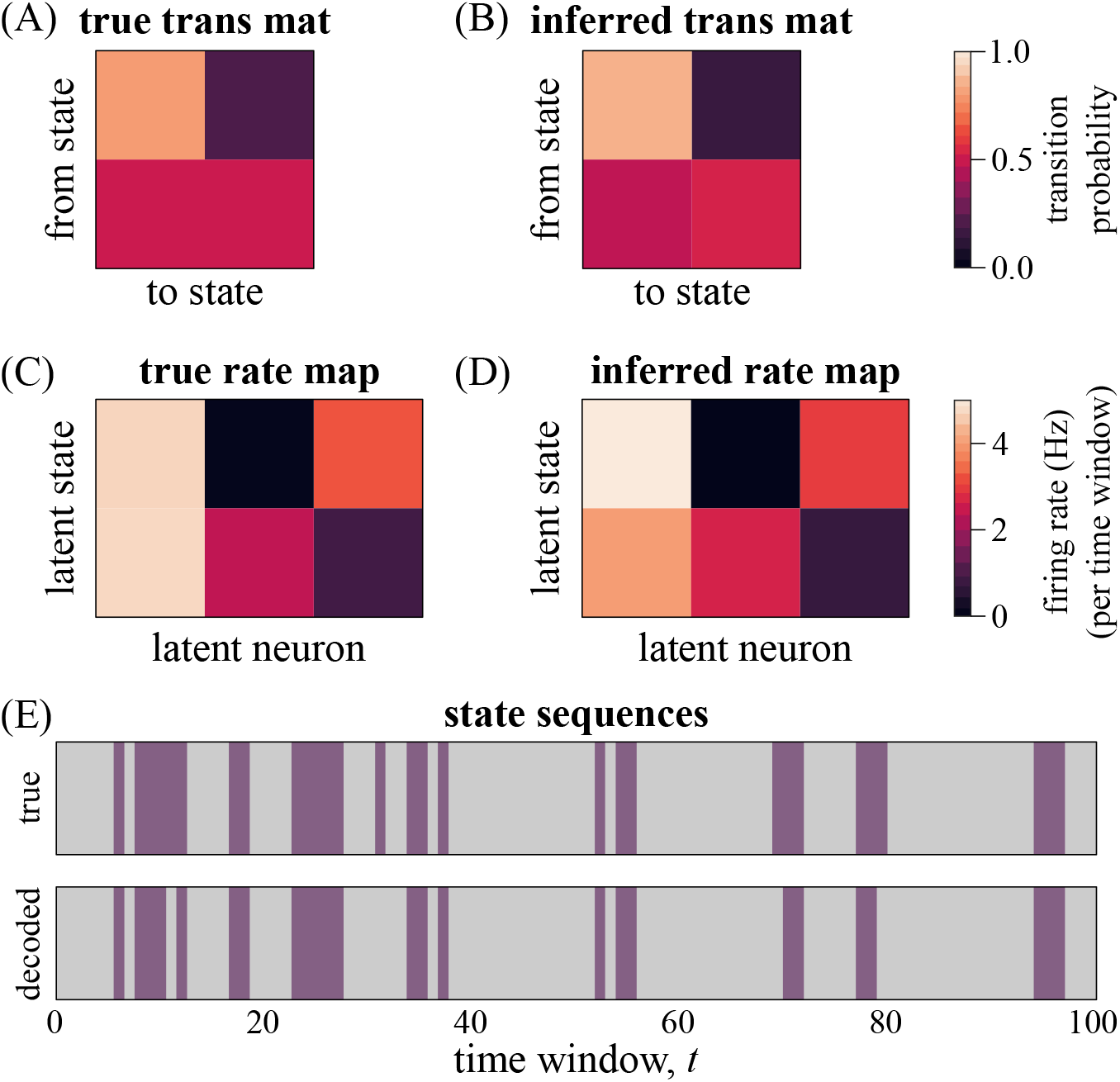
Parameter estimation and decoding example from synthetic dataset. (A) The true, and (B) inferred state transition probability matrices. (C) The true rate map that was used to generate the marks, and (D) the inferred rate map obtained by fitting the model. (E) Top: true state sequence generated from the transition matrix in (A); bottom: decoded state sequence. Colors indicate different states: *S_t_* ∈ {*z*_1_, *z*_2_}.

Notice that the parameters seem to have been estimated reasonably well, at least qualitatively. We define the relative error between matrices 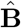 and **B** as

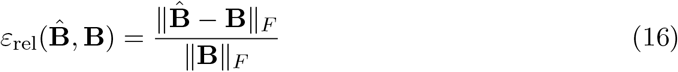

where ∥ · ∥_*F*_ is the Frobenius norm: 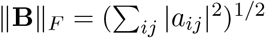.

The relative parameter estimation error for the state transition probability matrix was 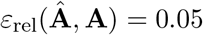, for the rate map, 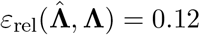, and the state decoding accuracy was 97.5%.

### 3.2 Misspecification of Hyperparameters

In the previous section we had assumed that both the number of latent states as well as the number of latent neurons were known, and the parameter estimation was performed with the exact numbers of each. In reality, we may neither know the “true” number of states, nor the number of neurons that participated in generating our data.

#### 3.2.1 Overspecification of Number of States

For the unknown number of states, a more advanced Bayesian treatment may be considered, where the number of states is itself learned directly from the data such as in the hierarchical Dirichlet process hidden Markov model (see e.g., Teh et al., 2006), or a likelihood-based approach may be followed, as described by Celeux and Durand (2008). However, as far as decoding accuracy is concerned, it may be argued that the HMM approach is largely insensitive to the precise number of states (see e.g., Maboudi et al., 2018, as well as the example in this section). In effect, choosing a larger number of hidden states partitions our latent space into a finer partition, and conversely, a lower number of states specifies a coarser partition. In an alternate view, if we consider a mapping from the state space (our domain) to some external space of interest (our codomain, e.g., an animal’s position in a maze), then we may expect to decode to the codomain reasonably well as long as we can find a surjective mapping; adding additional states should not affect our ability to find such a mapping, even though distinctness may become lost.

Indeed, we find that the decoding accuracy remains unchanged at 97.5% if we artificially use *Z* = 4 hidden states instead of the *Z* = 2 that were used to generate the data, and if we associate both dark and light purple states with “true” state *z*_1_, and both dark and light gray states with “true” state z_2_ (see Figure 5). This clearly illustrates the loss of distinctness, without affecting our ability to decode to the desired space (the original, data-generating state space in this instance).

**Figure 5:**
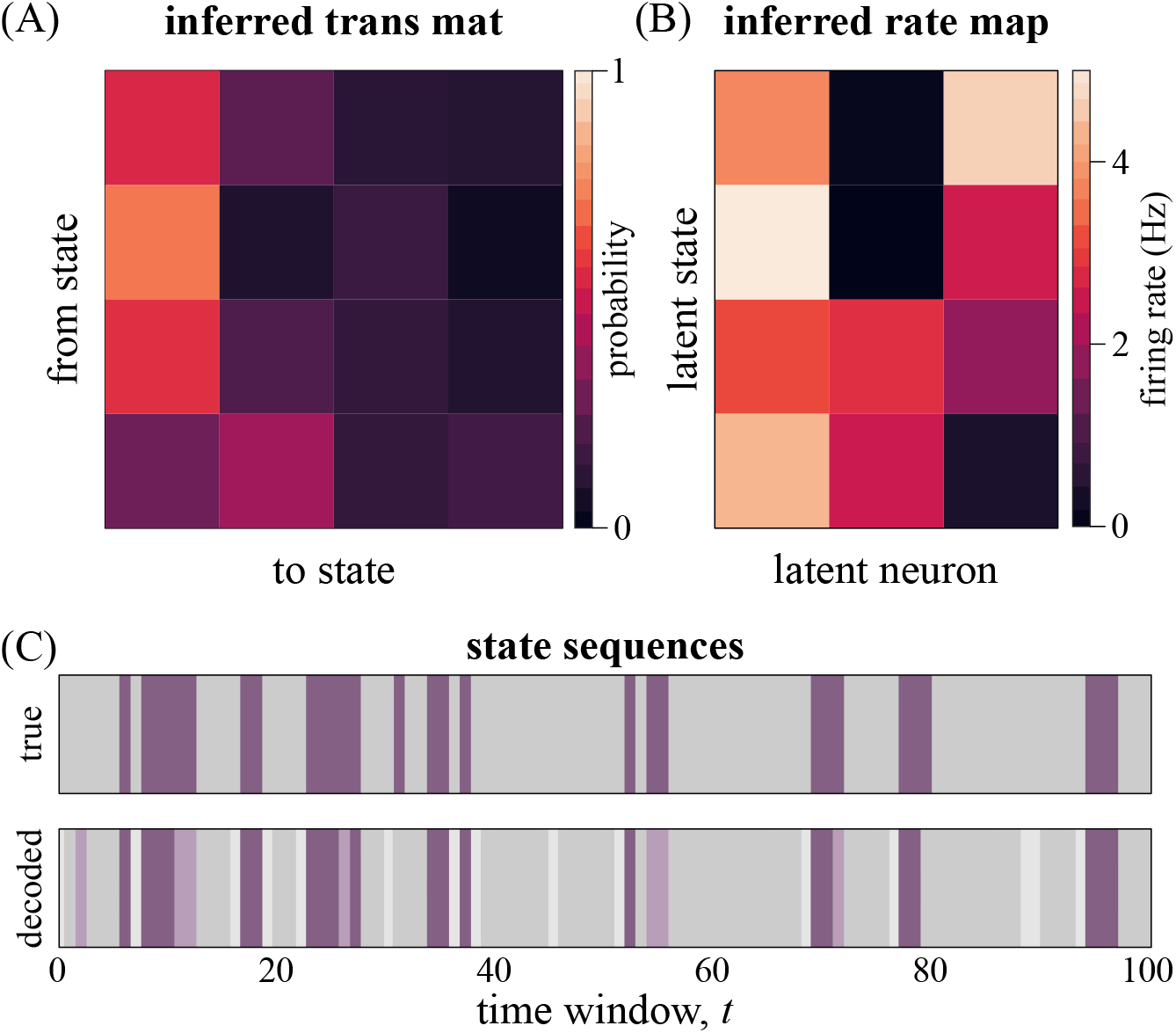
Overspecification of number of states. (A) The inferred transition matrix, and (B) the inferred rate map. (C) Decoding results. Top: true state sequence; bottom: decoded state sequence. Colors indicate different states: *S_t_* ∈ {*z*_1_, …, *z*_4_}.

#### 3.2.2 Overspecification of Number of Neurons

For the unknown number of neurons, things are a bit simpler. We may get some idea by simple visual inspection of our data, or by performing some reasonable clustering in our mark space. But as was the case for the number of hidden states, an overspecification of the number of neurons should have a relatively small effect on the model’s ability to decode (or to do most inference tasks, for that matter).

Indeed, if we over-specify *N* = 5 instead of the *N* = 3 neurons that were used to generate the data, we again find that our decoding accuracy is unchanged at 97.5% (see Figure 6). The relative estimation error in the transition matrix is also unchanged from when we had used the exact number of neurons.

**Figure 6:**
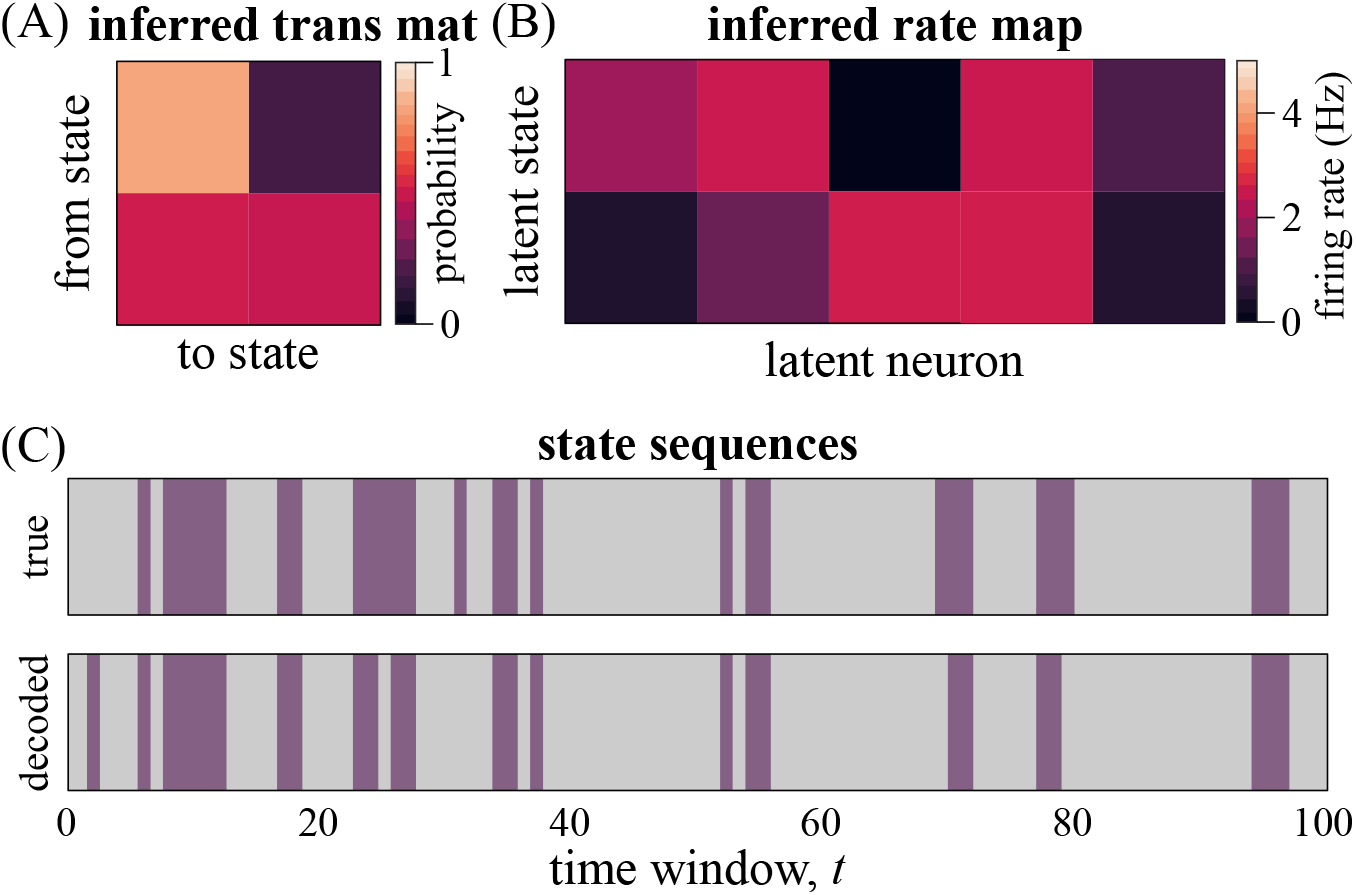
Overspecification of number of neurons. (A) The inferred transition matrix, and (B) the inferred rate map. (C) Decoding results. Top: true state sequence; bottom: decoded state sequence. Colors indicate different states: *S_t_* ∈ {*z*_1_, *z*_2_}.

This insensitivity to the number of neurons should not be very surprising, since neural decoding is often robust to subjective differences in the spike sorting process, where one researcher may combine two putative units together, while another may keep them as separate clusters. However, when we have limited data—such as during replay detection and analysis—the decoding robustness quickly fades, and small differences in spike sorting quality can have an outsized effect on analysis results. Thankfully, the clusterless approach presented in this paper (and in general) allows us to incorporate much more data than is typically available in a spike-sorting pipeline, thereby restoring some of the typical decoding robustness.

## 4. Application to Hippocampal Place Cell Data

We also fit our clusterless HMM to real neural data that were recorded from area CA1 in the rat hippocampus. In particular, a rat was trained to run back-and-forth on a 100 cm long linear track for liquid reward, and we subsequently recorded neural activity while the animal was performing this task. We expected that the clusterless HMM would capture the characteristic ensemble population “place cell” activity during periods in which the animal was running, and we expected to see that latent states correspond roughly to locations along the linear track (see e.g., Linderman et al., 2016; Maboudi et al., 2018).

### 4.1 Decoding Place Cell Activity with the Clusterless HMM

The animal was on the linear track for approximately 15 minutes. Periods of running activity were identified by applying a minimum speed threshold of 8 cm/s, and with this criterion, the animal was found to be running for a total duration of just over 4 minutes. Contiguous bouts of running (longer than 400 ms) were then partitioned into 400 ms observation windows. The distribution of these contiguous segments of running activity is shown in Figure 7.B, from where we can see that there are many short sequences (each consisting of only a few observation windows), and relatively few “long” sequences of 10 observation windows or more. With overwhelmingly short sequences, we can not expect the estimation of the transition probability matrix to be as reliable as if we had longer sequences, but we still find that the estimated transition probability matrix (Figure 7.C) is sparse, and strongly clustered around the diagonal and super-diagonal, with only a few other transitions scattered throughout. This sparse structure is suggestive of sequential hippocampal dynamics, as we would expect during running behavior (Maboudi et al., 2018).

**Figure 7:**
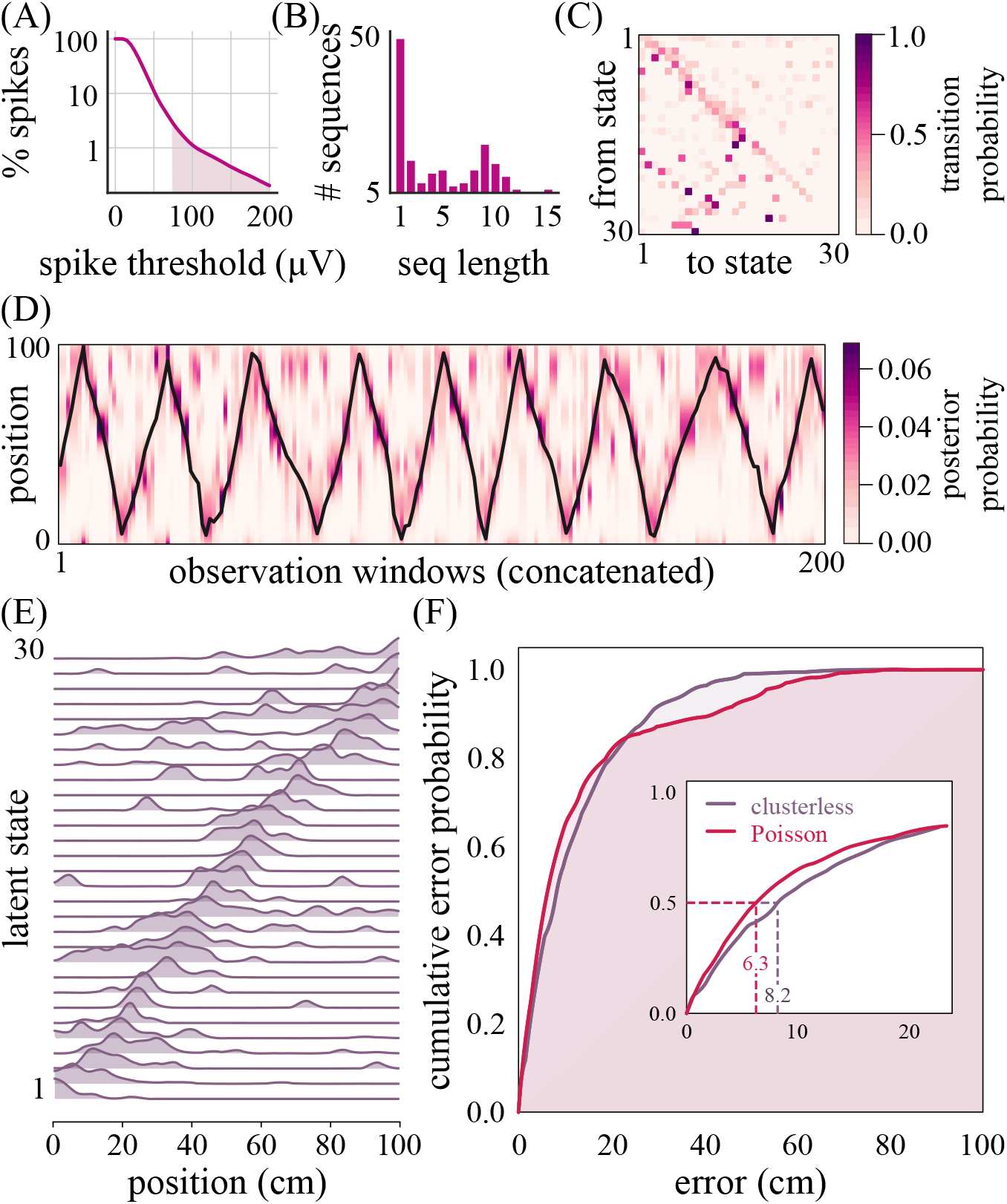
Clusterless hidden Markov model of hippocampal data during RUN. (A) Percentage of spikes retained as a function of detection threshold. We kept all spikes > 70 *μ*V (or ≈ 30%) (B) Histogram of sequence lengths (in number of observation windows). (C) Transition probability matrix. (D) Decoded posterior distribution over position for several run segments (200 × 400 ms windows). Black trace indicates animal’s actual position. (E) Latent state place fields (lsPFs) relating state space to animal position. (F) Cumulative distribution function of position decoding errors.

We included all (pyramidal and interneuron) spikes > 70 *μ*V for our clusterless HMM, which resulted in about twice as many spikes during run as compared to the manual-spike-sorted data (7,154 sorted vs. 14,453 unsorted spikes), and about 30% of the total number of spikes (using 0 *μ*V as baseline; see Figure 7.A).

By augmenting our dataset with some external correlate (the animal’s position in this case), we can learn a mapping between the state space and the external correlate. We refer to this mapping as the latent state place field (lsPF). In particular, we use our clusterless HMM to decode some neural activity to the the state space, and we use the associated position data to then learn a mapping between the state space and position. We notice that the states localize relatively well in space (Figure 7.E), and we can use these lsPFs to decode neural activity to position (Figure 7.D).

In Figure 7.D we see the posterior probability distribution over position for each 400 ms observation window, and more importantly, we see that the regions of highest probability typically coincide with the animal’s true position (black trace).

### 4.2 Comparison to Switching Poisson HMM

Unlike when we used simulated data, it is impossible to determine the “true” relative errors in estimating any of the parameters for the real neural data, especially since the latent states are merely mathematical abstractions (albeit with biological support or justification), so no true values for these parameters even exist. An alternative approach of evaluating our model’s performance is by analyzing its position-decoding ability instead. Here we have chosen to compare our clusterless HMM’s (position) decoding performance to the decoding performance of the switching Poisson HMM, where manually sorted data are used.

For both the clusterless and the switching Poisson HMMs, we kept as many of the hyperparameters the same as we could, including the observation window sizes, the sequences used for training and for testing, the number of (assumed) hidden states, etc. Using the manually-sorted spikes (where we have fewer spikes, but arguably higher quality spikes without interneurons or other “noise”), we obtain a median decoding error of only 6.3 cm, compared to a median decoding error of 8.2 cm for the clusterless approach (see Figure 7.F). However, the area under the curve (AUC) for the cumulative error distribution of the switching Poisson HMM is 87.3%, whereas the clusterless HMM achieved an AUC of 88.1%, although these differences did not appear to be statistically significant^2^.

Consequently, we may conclude that the clusterless HMM appears to have similar decoding accuracy compared to the switching Poisson HMM, even though no manual spike sorting was performed for the clusterless HMM.

## 5. Discussion and Future Directions

We have presented a new model, the clusterless HMM, which we have shown able to decode position just as well as the standard switching Poisson HMM, without access to the manually-sorted neuron identities. Even though the difference between the two approaches was not statistically significant here, others have shown that with the right dataset, the clusterless approaches can indeed improve decoding accuracy (Kloosterman et al., 2013; Matano and Ventura, 2018), and this potential improvement may be especially important for the study of severely data-limited analyses such as the study of replay (see e.g., Ackermann et al., 2017; Maboudi et al., 2018, for examples of applying the HMM framework to the identification and characterization of replay events).

In Section 2 we have only presented the sampling approach for spikes from a single (potentially multi-channel) probe. However, the extension to multiple probes is straightforward, and such an approach was applied to the real data example in Section 4. The key assumption for the multiprobe case is that each probe records from an independent subset of neurons, so that sampling can be restricted to neurons associated with each individual probe. Since probes are assumed independent, likelihoods can be computed by taking the product of the likelihoods for individual probes. The multiprobe extension therefore demands a sightly more sophisticated implementation, rather than any real algorithmic changes.

It is also possible to derive update and estimation equations for the mark distribution parameters directly in the Baum-Welch context (instead of using a separate estimation algorithm like the GMM that we have used in Sections 3 and 4), but such an approach would only really be beneficial if we assume that the mark distributions are state dependent.

One interesting consequence of our sampling approach to approximate *P*(**y**_*t*_ | *S_t_*) is that the data likelihood is no longer guaranteed to increase during each iteration of the EM algorithm (unless approximations are reused as suggested in Algorithm 1 in Appendix A). When we have found that the likelihood decreased during an iteration, we did not update the model parameters. Instead, we would repeat the iteration with a new set of samples, and if we again did not find an improvement in the likelihood, we would terminate the EM process. In practice, the early iterations made the biggest differences to the final likelihood, and those early iterations never seemed to result in a decrease in likelihood, so that whether we were to terminate immediately upon seeing a decrease in likelihood or whether we were to do some smarter iterative approach, it didn’t really seem to significantly affect the final results or model parameters.

Even though our clusterless HMM was able to recover the approximate parameters for the simulated dataset, and even though it clearly recovered temporal and task-relevant structure in the real dataset, there are a number of remaining challenges, the most important of which may be the high computational cost of the sampling approach that we have taken here. The model took about 6 hours to fit to the real dataset used here, which is already a relatively small dataset by today’s standards. However, several improvements could be made to speed up the computation time. For example, the samples in (6) are assumed to be independent, and so their calculation can be trivially parallelized. Some of the samples can also potentially be re-used in the forward backward algorithm (for example, the calculation of *α_j_*(*t* + 1) and *β_j_*(*t*) both make use of *P*(**y**_*t*+1_ | *S_t_* = *j*); see the Appendix). It is however fairly challenging to determine how many samples we need for sufficiently accurate estimates of *P*(**y**_*t*_ | *S_t_*), since it depends on many factors, including the degree of separability of the neuron clusters, the dimensionality of the feature space, the underlying firing rates of the neurons, and so on. Deriving an alternative way to approximate *P*(**y**_*t*_ | *S_t_*), or establishing some bounds or guidelines to determine the number of samples needed, is certainly a worthwhile future endeavor. Nevertheless, even with the relatively slow computation time, the clusterless HMM already provides a unique approach to the clusterless analysis of replay or other events where behavioral correlates are not available (e.g., during sleep), necessitating the use of unsupervised approaches.

## Acknowledgments

EA would like to acknowledge the helpful discussions with E. Buchanan, K. Kay, and X. Deng during the conception of this project.

## Animal Use

One male Long Evans rat (Charles River Laboratories) was implanted with a micro-drive array with independently adjustable tetrodes (implant coordinates3.66 mm AP and 2.4 mm ML relative to bregma). Tetrodes were slowly lowered into hippocampal CA1 over a period of about one week. Tetrode locations were verified by characteristic LFP waveforms attributed to the target area in conjunction with estimated tetrode depth. All experiments were approved by the Rice University Institutional Animal Care and Use Committee’s guidelines and adhered to National Institute of Health guidelines.

## Code Availability

A modified version of the Python package `hmmlearn` (https://github.com/hmmlearn/hmmlearn) has been made available at https://github.com/nelpy/hmmlearn. This modified package implements the multiprobe parameter estimation and decoding of our clusterless HMM with multivariate normal mark distributions.

## Appendix A: Parameter Estimation Algorithm

The iterative Baum-Welch parameter estimation procedure for our clusterless HMM is given in Algorithm 1. Algorithm 1 makes use of Algorithm 3 to approximate *P*(**Y**_*t*_ | *S_t_*), and Algorithm 2 to calculate 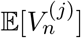.

### Algorithm 1 Clusterless HMM parameter estimation.

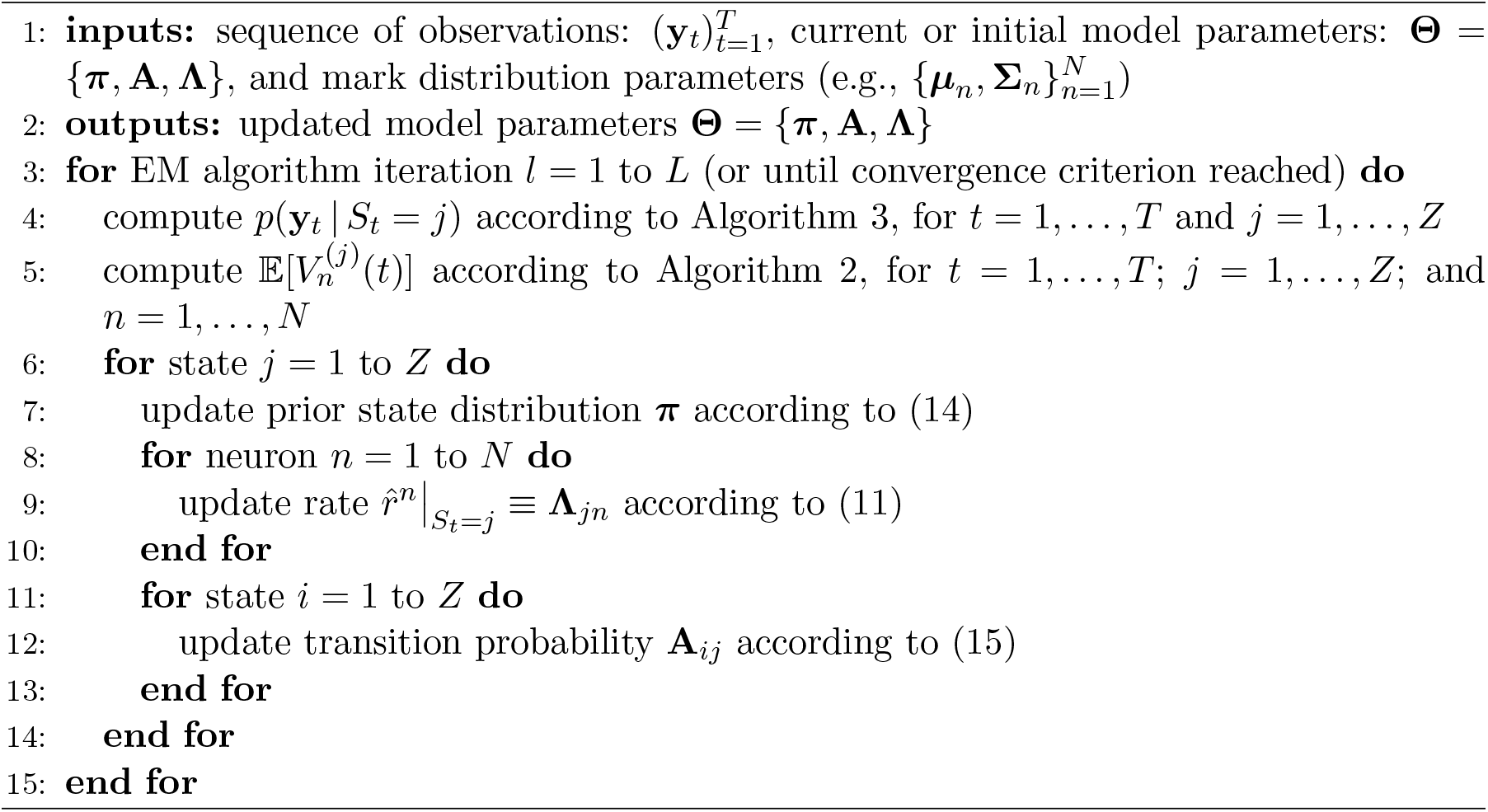

### Algorithm 2 Calculation of 𝔼[*V_n_^(j)^*].

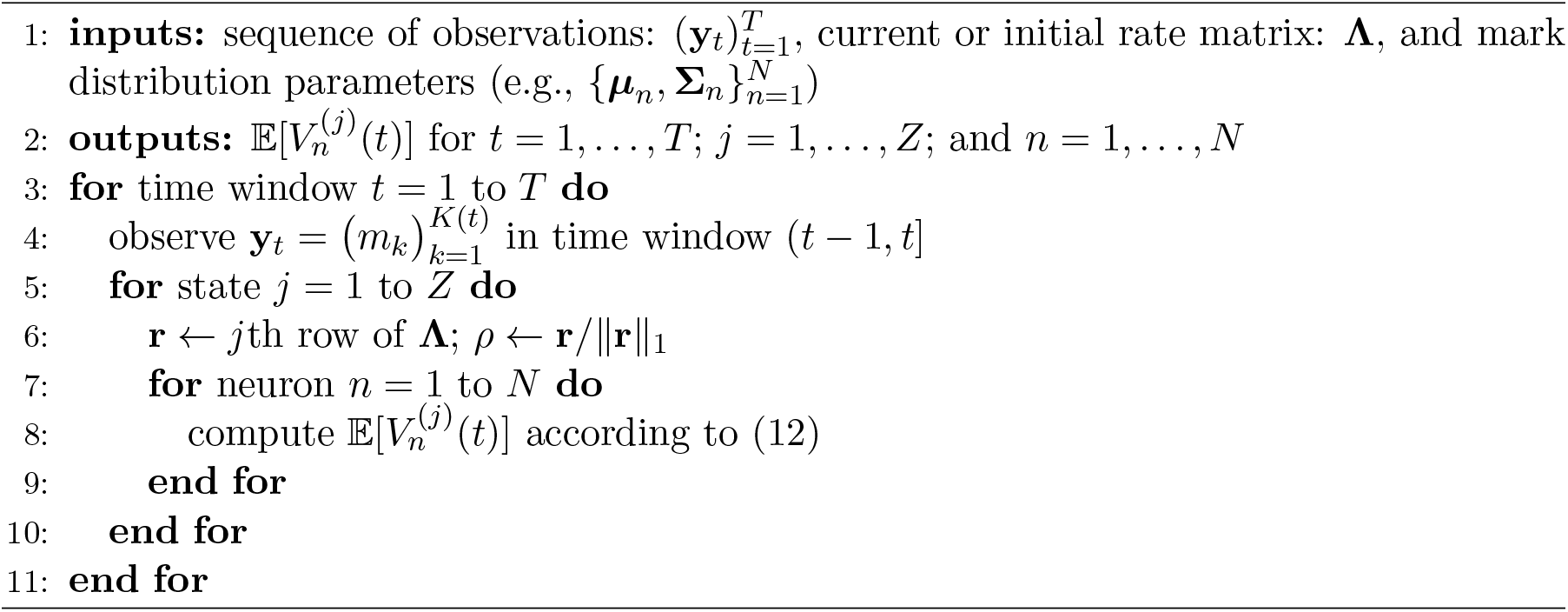

### Algorithm 3 Sampling approach to approximate *p*(**y***_t_* | *S_t_*).

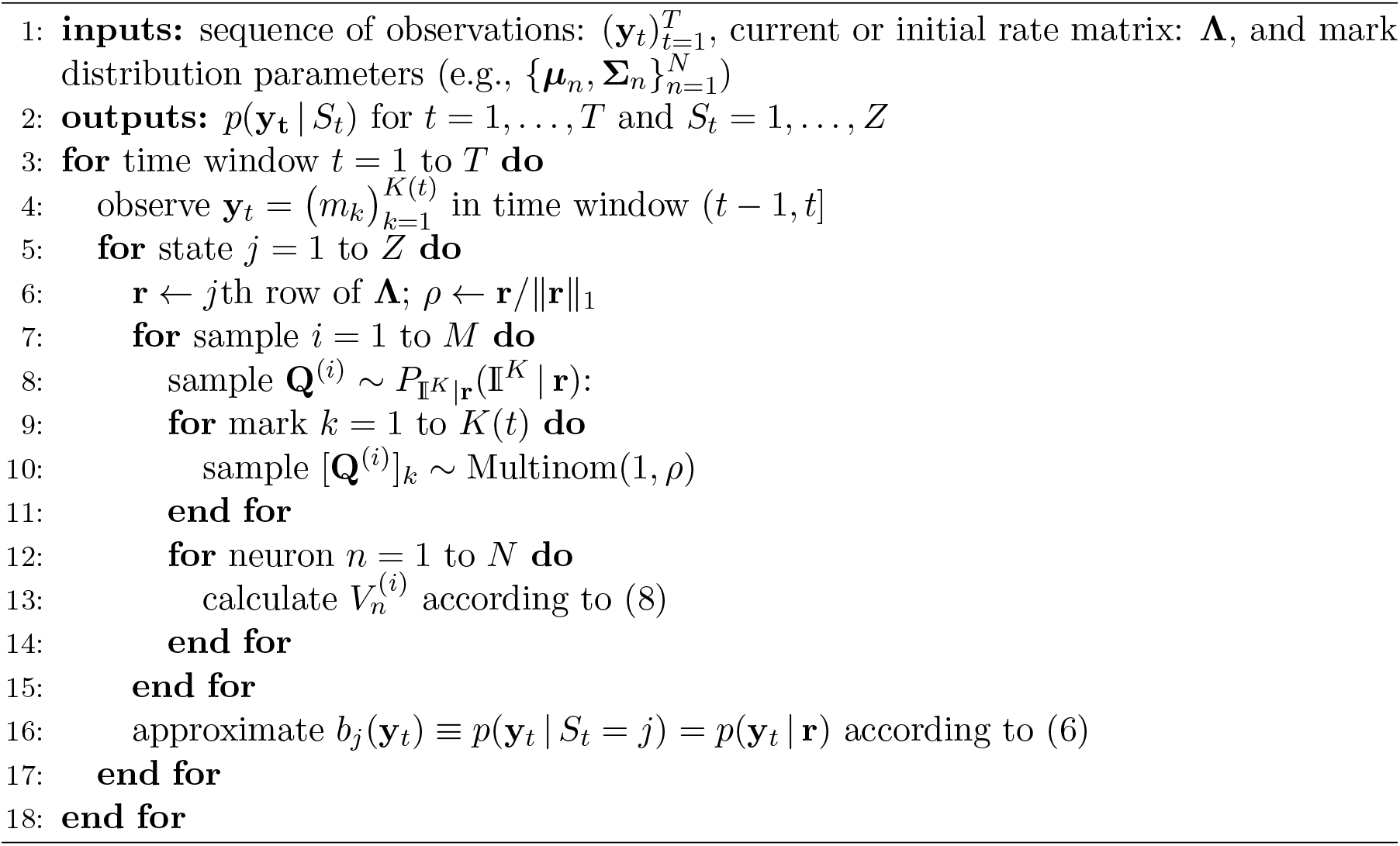

## Appendix B: Baum–Welch Algorithm

The Baum–Welch algorithm makes use of the forward-backward algorithm to compute the posterior marginals of all hidden state variables given a sequence of observations (Bilmes et al., 1998). That is, it computes *P*(*S_t_* | **y**_1:*T*_).

Following standard notation (and for notational simplicity), we define *b_j_*(**y**_*t*_) ≡ *p*(**y**_*t*_ | *S_t_* = *j*), which we evaluate by sampling as described in Section 2.3.

The forward algorithm gives us a way to compute the probability of seeing the partial sequence **y**_1_, …, **y**_*t*_ and ending up in state *i* at time *t*. That is,

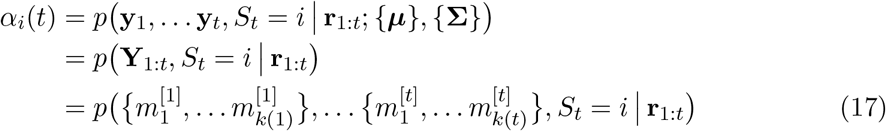

where **y**_*t*_ denotes the *t*th observation window, and where 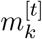 denotes the *k*th mark in time window *t*. In particular, *α_i_*(*t*) may be recursively computed as follows:

1. *α_i_*(1) = π_*i*_*b_i_*(**y**_1_)
2. 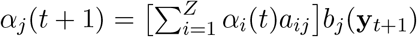
3. 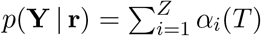

The backward procedure is similarly used to compute

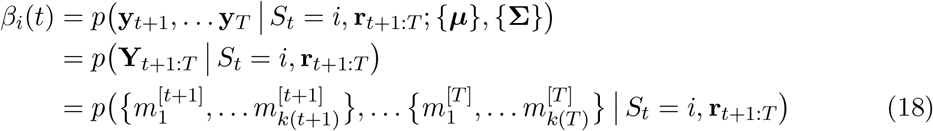

which is the probability of observing the partial sequence **y**_*t*+1_ … **y**_*T*_ given that we were in state *i* at time *t*.

Similar to before, *β_i_*(*t*) may be recursively computed as follows:

1. *β_i_*(*T*) = 1
2. 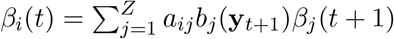
3. 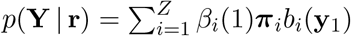

Having defined the forward and backward passes, we may now turn our attention to the parameter estimation problem more directly.

In particular, we define

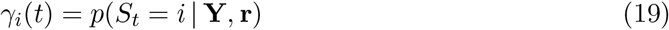

which can be shown (by Markovian conditional independence) to be equal to

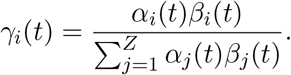

Let

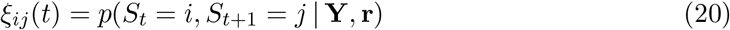

which we can show to be equal to

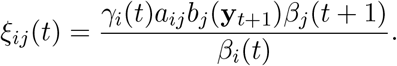

The parameter estimation updates then follow naturally in terms of *γ_i_* and *ξ_ij_* as described in Section 2.4. More specifically, the expected relative frequency spent in state *i* at time *t* = 1 gives us an estimate for **π**:

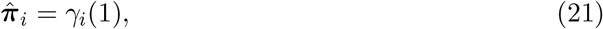

while the rate estimates are updated according to

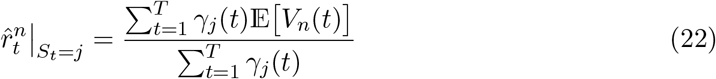

and the transition probabilities are updated according to

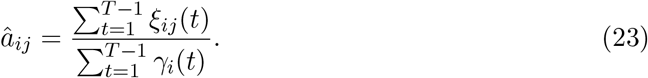

The update equations for multiple sequences of observations are easy to derive (see e.g., Rabiner, 1989, for details).

## Footnotes

1 Technically we may also need to estimate the number of hidden states, the number of hidden neurons, as well as the neuron cluster parameters, but those are typically estimated *a priori* and / or outside of the Baum-Welch algorithm, so that we do not consider them as part of the HMM parameter estimation process.

2 Statistical significance was evaluated with a bootstrap approach where we aggregated the AUCs of 5000 sampled CDFs (5000 for the switching Poisson group, and 5000 for the clusterless group), where each sampled CDF was obtained by randomly sampling, with replacement, from the pooled switching Poisson and clusterless HMM CDFs. Testing for a difference in the means of the two groups with Welch’s *t*-test resulted in a two-sided p-value of 0.37.

